# The distribution of *Apis laboriosa* revisited: range extensions and biogeographic affinities

**DOI:** 10.1101/2024.04.04.588077

**Authors:** Gard W. Otis, Man-Juan Huang, Nyaton Kitnya, Umer Ayyaz Aslam Sheikh, Abu ul Hassan Faiz, Chinh H. Phung, Natapot Warrit, Yan-Qiong Peng, Xin Zhou, Hliang Min Oo, Namoona Acharya, Kedar Devkota

**Affiliations:** School of Environmental Sciences, University of Guelph, Guelph, ON, Canada; Institute of Bee Health, Vetsuisse Faculty, University of Bern and Agroscope, Bern, Switzerland; CAS Key Laboratory of Tropical Forest Ecology, Xishuangbanna Tropical Botanical Garden, Chinese Academy of Sciences, Mengla, PR China; Department of Zoology, Himalayan University, Itanagar, Arunachal Pradesh, India; Trivedi Schools of Biosciences, Ashoka University, Sonipat, Haryana, India; Department of Entomology, University of Poonch Rawalakot, Rawalakot, AJK-Pakistan; Department of Zoology, Women University of Azad Jammu and Kashmir, Bagh, AJK-Pakistan; Mountain Bee Development JSC, 54/211 Alley, Khuong Trung St., Thanh Xuan District, Hanoi, Vietnam; Department of Biology and Center of Excellence in Entomology, Faculty of Science, Chulalongkorn University, Bangkok, Thailand; Department of Biomedical Science and Environmental Biology, Kaohsiung Medical University, Kaohsiung 80708, Taiwan; Department of Entomology, China Agricultural University, Beijing, PR China; University of Veterinary Science, Yezin, Nay Pyi Taw, Myanmar; Sub Metropolitan City Ward no. 12, Tulsipur, Dang District, Lumbini, Nepal; Faculty of Agriculture, Agriculture and Forestry University, Rampur, Chitwan, Nepal

**Author notes:** **Correspondence:** Gard W. Otis.

**Keywords:** *Apis laboriosa*, *Megapis*, Pakistan, Myanmar, Thailand, ecoregion, range map

## Abstract

*Apis laboriosa*, the Himalayan giant honeybee, inhabits the foothills of the Himalaya and neighboring mountainous regions. Here we revise its distribution in light of recent reports and discoveries. The range now extends from longitude 105.9°E in Cao Bang, Vietnam, in the east to 74.4°E in the Pir Panjal Range of western Himalaya, a linear distance of 3300 km, with the most notable new localities in northeastern Vietnam, central Myanmar, northern Thailand, and AJK-Pakistan. The species generally occurs at lower elevations in the eastern part of its range than in Nepal, northern India, and the border region between India and Pakistan. Most but not all of the new localities are within the range predicted by species distribution modelling. We discuss the new localities that fall outside of the predicted range, the biotic characteristics of the terrestrial ecoregions in which the species occurs, and the remaining regions that may harbor this spectacular honey bee species.

## 1. Introduction

> “Distribution maps are among the most fundamental and historically informative data of any biogeographic study.”

Parenti and Ebach (2009), Comparative Biogeography

The Himalayan giant honey bee, *Apis laboriosa* Smith, 1871, has been documented in mountainous regions of Asia, from northern India to northern Vietnam (Kitnya et al., 2020; Huang et al., 2022). It is a significant pollinator and a major source of honey in high-elevation regions of Asia (Batra, 1996; Gupta, 2014; Gogoi et al., 2017). The species was first collected and described from Yunnan, China (Moore et al., 1871), following which it was largely ignored until Maa (1953) resurrected it as a species, *Megapis laboriosa*, and provided a taxonomic description and key for identification. It again faded into obscurity until Sakagami et al. (1980) provided details for a number of distinct characters that clearly distinguish it from its sister species of mainland Asia, *Apis dorsata* F., 1793 (e.g., in workers, the colour of thoracic hairs and gastral tergites, size of ocelli, ocellar platform, ocellocular distance, malar length, and number of sting barbs; see Fig. 1a, b). Recently, genetic and morphometric analyses from several sites of sympatry in Arunachal Pradesh, India, confirmed that *A. laboriosa* and *A. dorsata* are distinct species (Kitnya et al., 2022). Classical morphological taxonomy (Kitnya et al., submitted) and recent genetic analyses (e.g., Arias and Sheppard, 2005; Raffiudin and Crozier, 2007; Lo et al., 2010; Kitnya et al., 2022; Bhatta et al., submitted) also differentiate all populations of these two species.

**Figure 1.**
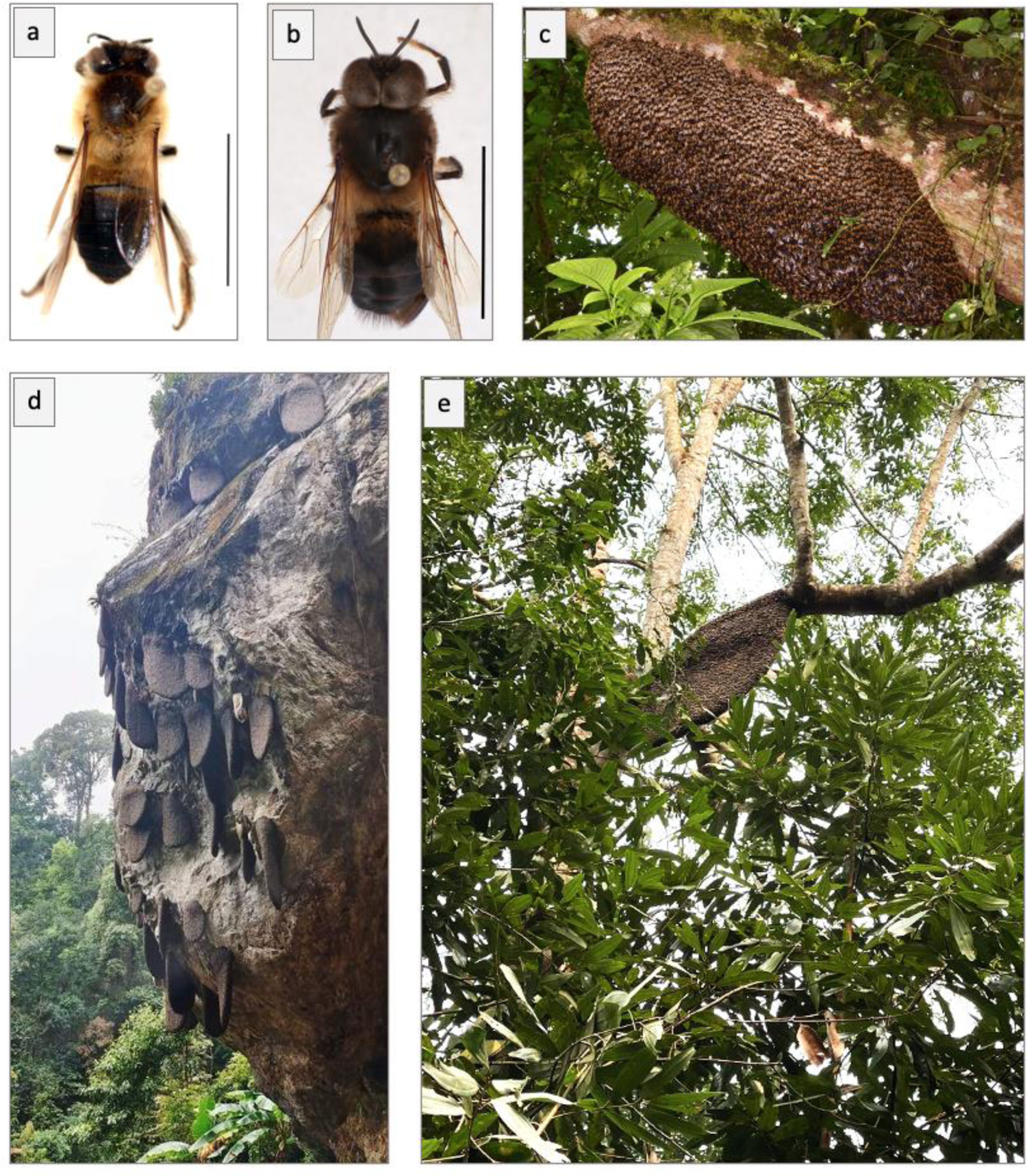
Habitus of worker (a) and drone (b) of *Apis laboriosa*; scale bar: 1 cm; specimens collected by B.A. Underwood, Kaski District, Nepal, 1675 m elevation (specimens in the Cornell University Insect Collection). (c) Wintering swarm (bivouc) on a tree branch 2–3 m above ground in Yen Bai Province, Vietnam, at 1200 m elevation; photo taken on 20 November 2022 by Eugene Popov. (d) Typical aggregation of nests on a cliff in Dien Bien Province, Vietnam; photo taken by Lo Van Anh of Son La Province. (e) *Apis laboriosa* nest constructed on a tree branch in Van Ban District, Lao Cai Province; photo taken by C. H. Phung.

Gradually, through accumulation of specimens, increasing amounts of research, and observations by naturalists, the distribution of *A. laboriosa* has been elucidated. Maa (1953) described its range as limited to “India (Sikkim; Assam); China (W. Yunnan). Probably also occurring in N. Burma.” Sakagami et al. (1980) created the first range maps for *A. laboriosa*, depicting it as a resident from central Nepal to Yunnan, China. Batra (1996) reported it from mountain valleys of Uttarakhand, India, ca. 600 km northwest of where it was known to occur in Nepal. Its distribution was further extended northward to Sichuan Province, China, and eastward to northern Laos (Otis, 1996) and at about the same time even further eastward to northwestern Vietnam (Trung et al., 1996). Gogoi et al. (2017) provided a generalized range map that showed it extending southward in the mountains near the border between northeastern India and western Myanmar. An updated distribution map that integrated information from the literature, iNaturalist reports, museum collections, and the authors’ personal observations added numerous localities in Bhutan, northeastern India, and northern Vietnam, and confirmed that it occurs in the Naga Hills of northeastern India and Chin Hills of west central Myanmar (Kitnya et al., 2020). Most recently, Huang et al. (2022) added several localities in China and Myanmar.

Kitnya et al. (2020) used rainfall and elevation patterns to suggest a number of regions where *A. laboriosa* was likely to occur but had not been documented: the Pir Panjal Range at the border between India and Pakistan; western Nepal; eastern Myanmar, and possibly as far south as Thailand; and along river valleys that extend into the eastern edge of the Hengduan Mountains of China. Subsequently, a species distribution model was generated for *A. laboriosa* by Huang et al. (2022). Species distribution modelling (SDM; also referred to by other names such as ecological niche modelling) allows one to predict the complete range of a species based on the environmental characteristics of sites where it has been documented to occur (Guisan et al., 2017). The range estimated by Huang et al. (2022) echoed most of the same regions predicted by Kitnya et al. (2020), as well as sites in northeastern Vietnam and a region on the Tibetan Plateau.

Here, we report several new discoveries that significantly extend the range of *Apis laboriosa* further to the east, south, and west. New discoveries in Xizang, China, provide hints about its possible migrations along river valleys. Evaluation of the biotic characteristics of the ecoregions (Olson et al., 2001; Dinerstein et al., 2017) and the climatic zones (Geiger, 1961; Kottek et al. 2006) it inhabits may be instructive in predicting other locations where this species does and does not occur.

## 2 Methods

To create our revised range map for *Apis laboriosa*, we began with a base map of Asia from OpenStreetMap-Boundaries (OSMBoundaries, 2024). Then, using QGIS (Version 3.32.2-Lima), we mapped locality records we obtained from various sources onto the base map. The lowermost layer consists of the records with coordinates reported in the supplementary data file to Kitnya et al. (2020). We have eliminated several of their records that lacked a source or for which the geographic coordinates had previously been approximated, resulting in 339 records, and we corrected one site in SE Nepal that previously had been entered incorrectly. We then added a layer showing the localities used by Huang et al. (2022) for species distribution modeling, excluding the records already reported by Kitnya et al. (2020) and those reported in GBIF (2023) and iNaturalist (2023a, b) (explained below).

In the third layer we have presented records obtained from the Global Biodiversity Information Facility (GBIF). GBIF (2023) reported occurrences of *A. laboriosa* in 8 datasets; we reviewed all of those (up to 3 November, 2023). Because records with coordinates from the Snow Entomological Museum, University of Kansas, and the US National Museum had been included by Kitnya et al. (2020) and most other GBIF records either lack coordinates or identifiable photos of the bees, we only obtained five new records from GBIF, all from the citizen science database Observation.org (Observation, 2023). Following the example of Dorey et al. (2023), we also checked the Symbiota Collections of Arthropods Network (SCAN, 2023); it contained only the records for specimens housed in the Snow Entomological Museum which already had been retrieved from the GBIF database. iDigBio (2023) lacked records for this species. Data for specimens housed in the Natural History Museum London were included by Kitnya et al. (2020). There were no specimens in the American Museum of Natural History, the Chicago Field Museum, or the Bishop Museum (Hawaii).

In the fourth layer, we have presented records from three citizen-science databases that contain identifiable images of bees in searches of *A. laboriosa* or *Indicator xanthanotus* Blyth, the yellow-rumped honeyguide that is intimately associated with nests of the bee: iNaturalist (iNaturalist, 2023a, 2023b; all posted records for the species checked between 28–30 October, 2023), and the Bhutan Biodiversity Portal (BBP, 2023) and Macaulay Library (2023) (sightings with photographs reported to eBird) (both reviewed on 7 November, 2023). Recent records that show abandoned honeycombs without identifiable bees were excluded. One additional locality was obtained from specimens housed in the insect collection of the Institute of Ecology and Biological Resources, Hanoi, Vietnam. After removing the records previously extracted from these websites and included by Kitnya et al. (2020), we found 61 verifiable postings.

We searched the Web of Science under the topic “Apis laboriosa” on 11 November, 2023 and retrieved several recently published papers. J.S. Xu (pers. comm.) confirmed the coordinates of the two locations from which mtDNA genomes were sequenced (Tang et al., 2003). Cao et al. (2023) reported five regions within Yunnan and Xizang, China, from which colonies were sampled; we have included two of those with narrow ranges of latitude and longitude. Gautam et al. (2022) studied pollination by honey bees at two sites in Uttarakhand, India. M. Kato (pers. comm.) confirmed one site where he and his associates observed *laboriosa* in Laos (Site S4 of Kato et al., 2020). Localities reported in Vietnam by Long et al. (2012) were not added because it seems that their identifications of *A. laboriosa* and *A. dorsata* were likely confused. These localities from literature sources were plotted in the fifth layer.

The 6th layer shows 39 new localities where *A. laboriosa* was observed by the authors or it was reported to them by reliable sources (e.g., honey-hunters) who included videos or photographs with identifiable images of bees.

Finally, we added the uppermost 7th layer with 16 observations that seem credible but are not supported by specimens, photographs, or videos.

To better understand the communities inhabited by *A. laboriosa*, we reviewed the ecoregions established by Olson et al. (2001) and Wikramanayake et al. (2002) and Köppen-Geiger climatic zones (Geiger, 1961; Kottek et al., 2006, Karki et al., 2016) that are known to be inhabited by *A. laboriosa*. The geographical terminology of Liu et al. (2022) for the Pan-Tibetan Highland region has been adopted.

## 3 Results

Figure 2 depicts the revised range of *Apis laboriosa*. It was created using data from refereed research publications, museum specimens, records with identifiable images of bees in publicly available databases, personal observations of the authors, and photos/videos and their coordinates submitted to the authors by honey-hunters and beekeepers. This map extends the distribution in three cardinal directions—east, south, and west—and fills in several previous gaps.

**Figure 2.**
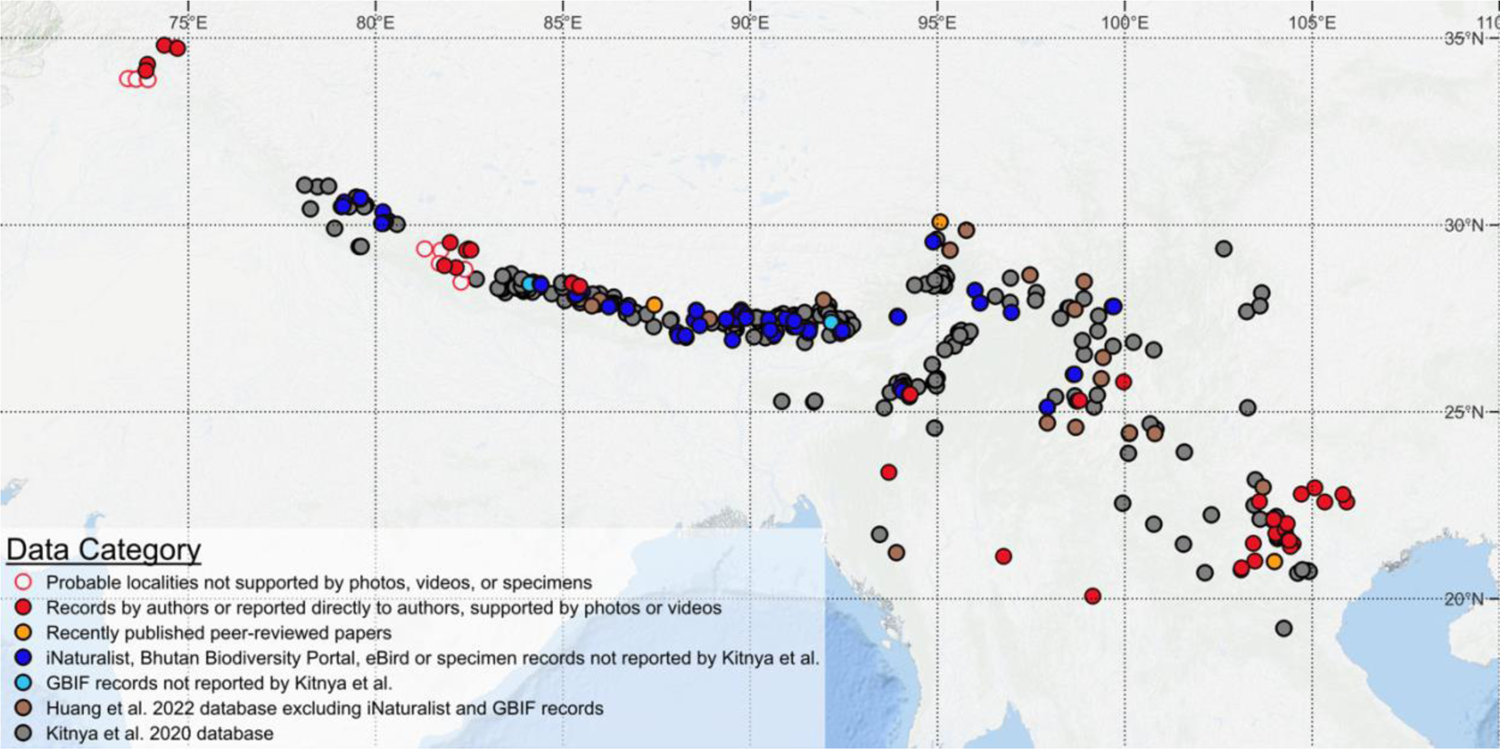
Revised distribution of *Apis laboriosa*. The colors of the dots reflect the source of locality information as indicated in the key at the lower left. Solid circles depict confirmed localities; open circles indicate probable localities based on credible reports that lack confirming photos, videos, or specimens.

In the east, in Vietnam and eastern Laos, the number of known localities was more than doubled, with the most easterly nests documented to date in Cao Bang Province, Vietnam (22.6°N, 105.9°E; Pham et al., 2023), nearly 200 km east of localities reported by Kitnya et al. (2020). Interestingly, although most nests in Vietnam are aggregated on cliffs (Fig. 1D), in some regions a higher proportion of nests are constructed on tree branches (Fig. 1E).

Across China, a number of new records add to the many already reported from there. Of note are the observations along several river valleys that extend northward into Xizang and Yunnan Provinces. For example, X. Zhou and his students recently collected specimens from several sites in Jilong County, Xizang, near the border of Nepal, and within the Kirong Tsangpo (called the Trishuli River in Nepal) drainage. Two nearly 50-year-old specimens in the National Zoological Museum of China were also collected in this same region. Cao et al. (2023) collected samples from several villages in Rikaze, Xizang, in the watershed of Phung Chu (the upper reaches of the Arun River) that extends northward from eastern Nepal. Further to the east, Huang et al. (2022) reported on specimens housed in the National Zoological Museum of China collected in the upper regions the Brahmaputra River: one along the Yarlung Zangbo River (in Motuozhen, Medog County), and another near Bomé along the Palong Zhangbo River, a tributary to the Yarlung Zhangbo, both in Nyingche Prefecture, Xizang. Cao et al. (2023) analyzed specimens collected in the nearby watershed of the Palong Zhangbo River. In Yunnan, Yang et al. (2015) reported the species from Deqin County (Shengpingzhen), near the Lancang River (upper Mekong River; included in Huang et al., 2022). We did not learn of any recent records from Sichuan Province.

In India, seven new records from Sikkim and West Bengal States (iNaturalist, 2023a, b) confirm previous old specimens collected in 1924 and 1938 in that region (Kitnya et al., 2020).

We have confirmed *A. laboriosa* in four districts of Karnali Province in western Nepal.

We also learned from agricultural extension agents, honey sellers, and honey hunters of additional three regions that we were unable to verify (e.g., Kalikot, Salyan and West Rukum Districts, Karnali Prov.; Bajura District, Sudurpashchim Prov.); these have depicted with open circles in Figure 2.

Several remarkable new records come from the southern and western edges of the range of the species. In extreme northwestern Thailand, *A. laboriosa* was independently observed foraging and nesting at the highest point in Doi Pha Hom Pok National Park, by nature photographers (iNaturalist, 2023a) and park personnel (Vorahab et al., 2024). In central Myanmar, bees collecting fluids from soil were photographed northwest of Taunggyi, in central Myanmar (21.1°N, 96.8°E), near mountains that exceed 2000 m in elevation at the western edge of the Shan Plateau. Y.Q. Peng collected the species above 3000 m on Mt. Victoria (Natma Taung), Chin State, western Myanmar (21.2°N, 93.9°E) (Huang et al., 2022).

Most surprisingly, we have confirmed that *A. laboriosa* inhabits the Neelum Valley, a region in northern Azad Jammu and Kashmir, Pakistan (AJK-P) dominated by coniferous trees (latitude 34.8°N). Active colonies were observed in Taobat at 2750 m elevation and foragers were observed on red and white clover (*Trifolium* spp.) in both Taobat and Arang Kel.

Aggregations of nests or foragers that were likely but not confirmed to be *A. laboriosa* were observed in Leepa; along the Neelum Jhelum River, and in Tolipir National Park. The giant honey bees reported by Khan et al. (2014) from Murree, Pakistan, at ca. 2000 m, were likely *A. laboriosa* that were incorrectly reported as *A. dorsata*. We have included these probable localities as open circles on Figure 2 (see also Supplementary Data File).

Our review of the ecoregions of Asia (Wikramanayake et al., 2002) indicates that *A. laboriosa* predominantly inhabits the Tropical and Subtropical Moist Broadleaf Forests Biome in several ecoregions (Northern Indochina Subtropical Forests, Chin Hills-Arakan Yoma Montane Rain Forests, Meghalaya Subtropical Forests, Eastern Himalayan Broadleaf Forests, Himalayan Subtropical Broadleaf Forests, Western Himalayan Broadleaf Forests). All of these ecoregions exhibit strong Himalayan biogeographic affinities (e.g., a predominance of oaks and numerous rhododendrons) and have moist to wet climates. Additionally, some localities in the western part of its range lie within the Himalayan Subtropical Pine Forests ecoregion and possibly Western Himalayan Subalpine Conifer Forests (e.g., at Rara Lake, Nepal and the Neelum Valley, AJK-Pakistan).

Over most of its range, from Yunnan, China, to Uttarakhand, India, and southward along the Arakan Mountains, *A. laboriosa* inhabits the Cwb (warm temperate, winter dry, warm summer) climate zone (Kottek et al., 2006). In contrast, in northern Vietnam and Laos the occupied climate zone is Cwa (warm temperate, dry winter, hot summer) and Cfa (warm temperate, fully humid, hot summer). In most parts of AJK-Pakistan, the inhabited regions are classified as Cfb (warm temperate, fully humid, warm summer), but the Neelum Valley is classified as Dfb (snow, fully humid, warm summer) (Geiger, 1961). These classifications are averages for the regions inhabited by the bee; in mountainous areas, microclimates can differ considerably over short distances due to local differences in elevation and rainfall.

## 4 Discussion

Our compilation of records provides the most up-to-date and comprehensive database for occurrences of *Apis laboriosa* (Supplementary Data File). We have indicated the categories of the sources of information, should others want to exclude some types of data from future analyses. The new observational records are conservative, in that for us to include them they were either made by one of the authors who is familiar with the species or were accompanied by a photo or video with identifiable bees, specimens, or publicly accessible DNA sequences.

We report here many new localities for *A. laboriosa* in northern Vietnam and Laos, more than one quarter of which were at elevations <1000 m, including what appears to be a combless wintering swarm (Fig. 1c; see Underwood, 1990), (Fig. 2). This species is widespread in the highlands of northern Vietnam, where we report for the first time that solitary nests are regularly constructed on tree branches, sometimes more frequently than on cliffs (C.H. Phung, unpublished data; Fig. 1e). Laos remains poorly sampled. The new records we report coupled with predictions from species distribution modelling (Huang et al., 2022) suggest that this species is likely much more widely distributed in northern Laos than has been documented. The mountains of northern Vietnam and Laos are the southeasternmost extension of the Himalaya Range (Sterling et al., 2006) and exhibit considerable Himalayan biogeographic affinities (e.g., Spitzer et al., 1993; Bain and Truong, 2004; Sterling et al., 2006; Bakalin et al., 2018, 2023) that undoubtedly influence the success of the bee there. In contrast, further south in the Central Highland region of Vietnam, despite some Himalayan influences in the flora (e.g., Vuong and Sridith, 2016; Wu et al., 2023), several searches by C.H. Phung and his colleagues for *A. laboriosa* in the region south of 15.5°N latitude and at elevations >1000 m have failed to detect this species. This may be a consequence of the habitats having unsuitable flora and/or climate, or a lack of connectivity with populations in the northern highlands resulting from the broad region between 16.5°–18.7°N latitude where elevations do not exceed 700 m.

We have now confirmed the species from several sites in western Nepal. An apparent gap in its distribution in that part of Nepal was discussed by Kitnya et al. (2020). Considering the new records (both confirmed and tentative), that gap likely represented a lack of scientific exploration in that remote part of the country. It is likely that this species occurs along the entire southern edge of the Himalaya in Nepal in a band of subtropical broadleaf forests (Wikramanayake et al., 2002) with Cwb climate (Karki et al., 2016), with extensions into landscapes dominated by conifers. The relatively dry environment of western Nepal may cause its occurrence there to be patchy (Karki et al., 2016). Additional confirmations through well documented observations and collections are warranted.

Both Kitnya et al. (2020) and Huang et al. (2022) predicted the occurrence of *A. laboriosa* along river valleys that extend into the Tibetan Plateau and Hengduan Mountains of China. This prediction is supported by observations of bees in the upper watersheds of the Trishuli River/Kirong Tsangpo, Arun River/Phung Chu, Brahmaputra River/Yarlung Zangbo/Tsangpo, and the Mekong River/Lancang. Despite being surrounded by high mountains, valleys, lower elevations along these rivers experience a subtropical climate dominated by broadleaf forests (Ni, 2000). *A. laboriosa* likely also occurs along the Salween River/Nu Jiang, the Nu River/Jinsha Jiang (the western tributary of the Yangtze River), and possibly more easterly tributaries of the Yangtze (i.e., Dadu, Min, and Jialing). It is not clear if it is a permanent inhabitant of these river valleys or if swarms migrate into them seasonally, as suggested by Underwood (1990) and Kitnya et al. (2020). Research on seasonal altitudinal migrations of this species is long overdue.

In Myanmar, *A. laboriosa* occurs near the summit of Natma Taung (Mt. Victoria), within the Chin Hills-Arakan Yoma Montane Rain Forests ecoregion (Wikramanayake et al., 2002). Cloud forests above 2000 m on the mountain are dominated by Himalayan tree taxa and have a distinctly Palearctic temperate flora (Wikramanayake et al., 2002). The Purvanchal and Arakan mountain ranges extend continuously from the Himalaya southward through northeastern India and western Myanmar, providing contiguous habitat and a corridor for bee dispersal along the N/S axis of the mountains. Considering all these factors, the presence of *A. laboriosa* on Natma Taung was anticipated.

Discoveries of *A. laboriosa* in central Myanmar and northwestern Thailand extend its distribution southward. These localities are far (ca. 350 km and 240 km respectively) from the closest known populations in Yunnan, China. It was predicted that the species may occur on Doi Pha Hom Pok, Thailand (Kitnya et al., 2020; Huang et al., 2022), as has now been confirmed (iNaturalist 2023a; Voraphab et al., in press). On the Shan Plateau of eastern Myanmar, relatively large areas exceed 1000 m in elevation (see map provided by Evers and Taft 2016), the approximate elevation at which the species was photographed in Pindaya Township, Shan State, and where the climate is classified as “temperate, dry winter, hot summer” (Cwa) (Geiger 1961; Kottek et al. 2006). Scattered over this region at higher elevations, the climatic zone is “temperate, dry winter, warm summer (Cwb) and more suitable for the bee. *Apis laboriosa* likely inhabits many of these high elevation sites on the Shan Plateau, some of the more southerly of which were not predicted by Huang et al. 2022. It may even occur as far south as Nattaung (latitude 18.8°N), the highest mountain (2623 m) of the Karen Hills, and the high elevation region surrounding that mountain.

The most exciting new discoveries are located in the Pir Panjal Range, in regions administered by Pakistan, ca. 530–630 km northwest of the closest known populations of *A. laboriosa* in Uttarakhand, India. Visual observations in four different regions of AJK-Pakistan were made by U.A.A. Sheik and A.H. Faiz who are very familiar with *Apis dorsata*, the species with which it would most likely be confused. They identified *A. laboriosa* at four sites in the Neelum Valley, Leepa Valley, and along the Jhelum River on the basis of coloration of bees that were observed both foraging on flowers and nesting on rock cliffs. In they Neelum Valley they were observed in September at high elevations (e.g., >2500 m). Unfortunately, specimens and photographs that support these observations are currently lacking. The climate in the sites the bees have been reported from spans zones Cfb (Temperate, no dry season, warm summer), Dfb (cold, no dry season, warm summer), and Dsb (cold, dry summer, warm summer) in the upper Neelum Valley; Geiger 1961). Interestingly, some of these sites are dominated by conifers (e.g., Ecoregion 31: Himalayan subtropical pine forests) rather than broadleaf evergreen forests (Wikramanayake et al., 2002), suggestive that the niche of the species is broader than indicated by the predominantly broadleaf forests it usually inhabits elsewhere. Kitnya et al. (2020) predicted its occurrence in AJK-Pakistan. In contrast, although Huang et al. (2022) indicated it may occur near Srinigar, India, their species distribution modeling results did not include regions administered by Pakistan. Earlier reports of *Apis dorsata* at three sites at ca. 2000 m elevation near Murree, Pakistan, were likely of *A. laboriosa* (Khan et al., 2014; specimens not retained, K.A. Khan, pers. comm.). We have depicted unconfirmed sites with open circles on our revised distribution map (Fig. 2) in the hope that others will be stimulated to confirm the occurrence of this disjunct population and study its biology in the Pir Panjal Mountain Range.

Species distribution modelling predicted most of the new sites in which *A. laboriosa* has now been documented, showing the general utility of this research tool. However, it failed to detect the Central Myanmar and AJK-Pakistan localities. Conversely, it did predict its occurrence on the southwestern Tibetan Plateau (between ca. 81.0–85.3°E longitude; Huang et al., 2022), for example in the upper basin of the Yarlung Zangbo River in Zhongba and Saga Counties, Xizang. This is a semi-arid region dominated by grasslands, with mean annual temperature <0°C (Wang et al., 2020). The climate over most of this region has been classified as “Polar, tundra” (ET), with small portions of the region classified as “Snow, winter dry, warm summer (Dwb)” and “Snow, winter dry, cool summer” (Dwc) (Kottek et al., 2006). As discussed above, the species has been documented along the Yarlung Zangbo River, but much further to the east (Medog County, Nyingche/Linzhi) which, along with many other parts of Yunnan, has broadleaf forests and climate classified as “Warm temperate, winter dry, warm summer” (Cwb) (Kottek et al. 2006). The high elevation (>4000 m), cold climate, and alpine meadow vegetation of the Tibetan Plateau north of Himalaya (Li et al. 2010), as well as the lack of river valleys that could serve as migration corridors to warmer sites within migration distance on the southern edge of the mountains, make it questionable that *A. laboriosa* inhabits this region. Two sites on the Tibetan Plateau reported by Kitnya et al. (2020), the geographic coordinates of which were estimated from a general description of their location, have been removed from the current data (Supplementary Data). The environmental conditions at those sites may have influenced the species distribution modelling of Huang et al. (2022). The questionable existence of this species in this part of southwestern Xizang should be checked through field work.

Our analysis shows that *Apis laboriosa* occurs from northeastern Vietnam to AJK-Pakistan, an linear east-west distance of ca. 3300 km, and from Sichuan, China, in the north to northern Thailand in the south. It is a regular inhabitant of high elevation, moist evergreen broadleaf forests with strong Himalayan floral influences. In the western portion of its range it extends into coniferous forests. We anticipate that *A. laboriosa* will eventually be confirmed in appropriate habitats/climates of: (1) the mountains of Himachal Pradesh and (2) Jammu in regions administered by India; (3) the sites we have provisionally shown, as well as the Murree Hills and the Galis in regions administered by Pakistan; (4) additional sites in an elevational band with appropriate climate and habitat across western Nepal; (5) at numerous sites in northern Laos, particularly in the northeast; and (6) scattered over the Shan Hills of eastern Myanmar, potentially as far southward as the Karen Hills. We hope that this reassessment of the range of *Apis laboriosa*, coupled with hints of ecological differences over this large region, will inspire more detailed studies of its ecology, population genetics, migratory behavior along river systems, and role as a pollinator within agroecosystems (e.g., Gautam et al., 2022) and natural communities (e.g., Kato et al., 2020).

## Data Availability

All locality information used to generate the distribution map (Fig. 2) is presented in the Supplementary Data.

## Ethics Statement

Ethics review and approval were not required for this study. *Apis laboriosa* is not endangered, nor is it listed under CITES.

## Author Contributions

GWO and NK: conceptualization; MJH, YQP, UAAS, AHF, CHP, NW, XZ, HMO, NA: field work and contacting honey hunters and beekeepers; GWO: searching websites and literature for locality data; NK: creation of the distribution map; GWO: photos of the worker and drone; CHP: photo of tree nest; GWO: writing manuscript drafts. All authors edited and approved the manuscript.

## Acknowledgements

Gard Otis dedicates this paper to Makhdzir Mardan (1953–2022). Makhdzir first introduced Gard to *Apis dorsata* in Malaysia in 1986, a profound experience that stimulated his interest in Asian honey bees that continues to the present. Throughout his career, Makhdzir championed research on and conservation of giant honey bees. We are indebted to numerous honey-hunters and beekeepers who shared locations, photographs and videos of *A. laboriosa* nesting sites. Steve Paiero, University of Guelph, assisted with the photographs of worker and drone specimens. Eugene Popov kindly allowed us to use his photo of a wintering swarm posted to iNaturalist. Similarly, Lo Van Anh gave us his permission to include his photo of an aggregation of bee nests. Prof. Chaodong Zhu and his team (Zeqing Niu, Qingtao Wu) from the Institute of Zoology, Chinese Academy of Sciences led the collection trip in 2023 in Jilong, Xizang, China, and kindly shared sample information included in this paper. We thank Itsarapong Voraphab (Thailand Dept. of National Parks, Wildlife and Plant Conservation), and Chawatat Thanoosing, Nontawat Chatthanabun, Pakorn Nalinrachatakan, Prapun Traiyasut, and Chawakorn Kunsete (Chulalongkorn University, Bangkok) for their contributions to the discovery of *A. laboriosa* nests in northern Thailand. Bilal Abdulah, Umer Ghaffar and Wajahat conducted field work that confirmed the species in the Neelum Valley, AJK-Pakistan.

